# Characterization of extracellular vesicles isolated from *Sorghum bicolor* reveals a conservation between monocot and eudicot extracellular vesicle proteomes

**DOI:** 10.1101/2023.05.25.542161

**Authors:** Timothy Chaya, Aparajita Banerjee, Brian D. Rutter, Deji Adekanye, Jean Ross, Guobin Hu, Roger W. Innes, Jeffrey L. Caplan

**Affiliations:** Department of Plant and Soil Sciences, University of Delaware, Newark, Delaware 19716, USA; Department of Biology, Indiana University, Bloomington, Indiana 47405, USA; Brookhaven National Laboratory, Upton, New York 11973, USA

## Abstract

Plant extracellular vesicles (EVs) are membrane bound organelles involved mainly in intercellular communications and defense responses against pathogens. Recent studies have demonstrated the presence of proteins, nucleic acids including small RNAs, and lipids along with other metabolites in plant EVs. In this paper, we described the isolation and characterization of extracellular vesicles from *Sorghum bicolor*. Nanoparticle tracking analysis, dynamic light scattering, and cryo-electron tomography showed the presence of a heterogeneous population of EVs isolated from the apoplastic wash of sorghum leaves. Cryo-electron microscopy revealed that EVs had a median size of 110 nm and distinct populations of vesicles with single or multiple lipid bilayers and low or high amounts of contents. The heterogeneity was further supported by data showing that only a subset of EVs that were stained with a membrane dye, Potomac Gold, were also stained with the membrane-permeant esterase-dependent dye, Calcein-acetoxymethyl ester. Proteomic analysis identified 437 proteins that were enriched in multiple EV isolations, with the majority of these also being found in the EV proteome of Arabidopsis. These data suggest a partial conservation of EV contents and function between the monocot, sorghum, and a distantly related eudicot, Arabidopsis.

## Introduction

Extracellular vesicles (EVs) are lipid bilayer membrane-enclosed nanoparticles released by the cells of various organisms ranging from prokaryotes to eukaryotes into extracellular spaces (Yáñez-Mó et al., 2015; Tkach and Théry, 2016; Cui et al., 2020; LeClaire et al., 2021). EVs mediate intercellular communications, both proximal and distal to the place of origin, through the delivery of proteins, nucleic acids including small RNAs, lipids, and other metabolites (Yáñez-Mó et al., 2015; Tkach and Théry, 2016; Maas et al., 2017; Wu et al., 2019; Liu et al., 2020a; Woith et al., 2021). Previously, EVs were classified into three main categories: apoptotic bodies (1000-5000 nm in diameter) generated during programmed cell death via cell fragmentation and blebbing of plasma membrane, microvesicles (100-1000 nm in diameter) released by direct budding and shedding of plasma membrane, and exosomes (30-150 nm in diameter) originated intracellularly as multivesicular bodies (MVBs) that fuse with the PM to release intraluminal vesicles outside the cell (György et al., 2011; Théry, 2011; Pol et al., 2012; Akers et al., 2013; Raposo and Stoorvogel, 2013; Lawson et al., 2016; Rutter and Innes, 2017; Zempleni et al., 2017; Woith et al., 2019). However, lacking clear evidence of biogenesis or biomarkers, it is necessary to categorize EVs with physical characteristics such as buoyancy, molecular cargo or size (Théry et al 2018). In the field of EV research, mammalian EVs are widely studied. They are released from almost all cell types and body fluids and play critical roles in immune responses and the progression of various diseases (Colombo et al., 2014; Woith et al., 2019). Based on their significance as carriers of biomolecules as well as potential vehicles equipped to cross challenging pharmacological barriers, mammalian EVs have been extensively explored in clinical and therapeutic applications (Pol et al., 2012; Elsharkasy et al., 2020).

Compared to the comprehensive research encompassing mammalian EVs, the research in the field of plant EVs is still in its infancy. Plant EVs were first discovered in the 1960s, where they were described in carrot cell culture and in wheat (Manocha and Shaw, 1964; Halperin and Jensen, 1967). The current evidence suggests plant EVs play a role in intercellular and extracellular communications, cross-kingdom exchange of small RNAs, plant immune responses, and plant-microbe symbiosis (Regente et al., 2017; Rutter and Innes, 2017; Cai et al., 2018; Rutter and Innes, 2018; Cui et al., 2020). Despite these new findings, there remain many fundamental questions regarding plant EVs, even six decades after their first discovery. Further research needs to be done to shed light on their biogenesis, morphology, packaging of cargoes, mechanism of passage through the cell wall, inter-kingdom movement and EV cargo delivery to name a few.

Various biofluids including blood, urine, cerebrospinal fluid, lymphatics, tears, saliva and nasal secretions, ascites, and semen have been used as sources of mammalian EVs (Akers et al., 2013). Stepwise differential centrifugation is used most often to isolate EVs from mammalian biofluids. Briefly, fluids are processed at a low centrifugation speed to remove cellular debris and large particles followed by ultracentrifugation at a higher speed, typically at 100,000 x g, to pellet EVs (Théry et al., 2006; Akers et al., 2013; Gardiner et al., 2016). Further purification can be obtained by ultracentrifugation of the crude EVs using a discontinuous density gradient (Raposo et al., 1996; Akers et al., 2013).

In the plant sciences, EVs are most frequently isolated from apoplastic wash fluids. Such fluids are obtained by vacuum-infiltrating a buffer into leaves and using slow centrifugation speeds to push the buffer out of the leaves through stomatal openings. (Rutter and Innes, 2017; Liu et al., 2020a; Schlemmer et al., 2021). This process effectively rinses the extracellular spaces inside the leaves and allows researchers to collect any molecules or particles therein. To date, EVs from the wash fluids of Arabidopsis and barley leaves have been isolated using differential ultracentrifugation. Additionally, many studies have isolated plant-derived, edible nanoparticles (PDVs) from the juices of various fruits and vegetables, such as grapefruit, grape, lemon, strawberry, orange, ginseng, ginger, carrot, broccoli, and cabbage (Wang et al., 2013; Mu et al., 2014; Raimondo et al., 2015; Zhang et al., 2016; Deng et al., 2017; Yang et al., 2018; Cao et al., 2019; Iravani and Varma, 2019; Berger et al., 2020; Yang et al., 2020; Man et al., 2021; Perut et al., 2021; Savci et al., 2021; You et al., 2021). PDVs have potential therapeutic and clinical applications, but since they are often a mixture of vesicles with intracellular and extracellular origins, PDVs should not be used to study the natural formation and function of EVs.

While there is an understanding that these vesicles play a key role in the response to various pathogens, particularly fungi, defining their function in the immune response is an ongoing process. It is also unknown how conserved plant EVs are in cargo and composition from monocots to eudicots.

To expand our knowledge of plant EVs, we describe here methods for isolating and purifying EVs from the important cereal crop *Sorghum bicolor*. Vesicles from this monocot could be isolated from the wash fluids of leaves and were similar both in morphology and protein content to EVs from Arabidopsis. In this study, we described an optimized a protocol for EV isolation from sorghum leaves using differential centrifugation methods. We also characterized the isolated EVs using nanoparticle tracking analysis, dynamic light scattering analysis, and various microscopic techniques.

## Materials and Methods

### Plant material and growth conditions

*Sorghum bicolor* (BTX623) was used in this study. Seeds were potted using PRO-MIX BK55 growing medium. Plants were grown under 16 h days, 27 ⁰C, 350 µmol m^-2^ s^-1^, 65% humidity. Plants were grown for ∼14 days before harvesting for the isolation of extracellular vesicles.

### EV isolation protocol

EVs were isolated from ∼14 days old plants following the protocols used by Rutter et al. with some modifications (Rutter and Innes, 2017). Before harvesting, 0.5 cm of the leaf tips of each plant were cut to make the infiltration easier. The plants were harvested just above the soil and submerged in water to remove damaged plant material and residual soil. In small groups, the plants were prepared for vacuum infiltration in a 1L glass beaker using a glass petri dish lid on top of the plant tissue to keep the leaf material submerged in vesicle isolation buffer (VIB; 20 mM MES, 2 mM CaCl2, and 0.1 M NaCl, pH 6.0). The plants were vacuum infiltrated with VIB using five, 20 sec cycles at -0.1 MPa with reshuffling of the plants in between cycles. After removal of plants from the 1L flasks, the remaining buffer was saved for subsequent isolation of EVs that had been released into the buffer during the vacuum infiltration process. We refer to this as vacuum-based apoplastic wash (vAW). For collecting the-centrifugation based apoplastic wash (cAW), small batches of VIB-infiltrated plants were bundled into bouquets with a small strip of parafilm and placed stem side down inside 60 mL syringes. The 60 mL syringes were loaded into 250 mL centrifuge bottles (Nalgene) that were widened to fit a 60 mL syringe. The cAW was collected by spinning at 700 x g for 20 min at 4 ⁰C (JA-14 rotor, Beckman Coulter; Avanti J-20^TM^ centrifuge, Beckman Coulter). vAW was filtered through a single layer of Miracloth and was combined with the cAW. The combined cAW and vAW solutions were then centrifuged using 29 x 104 mm (50 mL) polypropylene centrifuge tubes (Beckman Coulter, # 357007) at 10,000 x g for 1 h at 4 ⁰C (JA-17 rotor, Beckman Coulter; Avanti J-20^TM^ centrifuge, Beckman Coulter). The supernatant from this step was decanted into the new tubes and centrifuged at 39,800 x g for 2 h at 4 ⁰C (JA-17 rotor, Beckman Coulter; Avanti J-20^TM^ centrifuge, Beckman Coulter). The supernatant was then decanted and discarded. The pellet from each tube was resuspended in chilled VIB and pooled together. The resulting solution was then centrifuged again at 39,800 x g for 2 h at 4 ⁰C (JA-17 rotor, Avanti J-20^TM^ centrifuge, Beckman Coulter). The supernatant was decanted, and the pellet (P40) was resuspended in 100 – 250 µl of 0.2 µm filtered 20 mM Tris-HCl pH 6.0.

### Density gradient purification

Crude sorghum EVs from the P40 were further purified using discontinuous iodixanol gradients (Optiprep, Sigma-Aldrich) following the protocol described by Innes and Rutter (Rutter and Innes, 2017). Briefly, 60% aqueous iodixanol stock solution was used to prepare 40% (v/v), 20% (v/v), 10% (v/v), and 5% (v/v) iodixanol dilutions in VIB. Next, the discontinuous density gradient column was prepared by carefully layering 3 mL of 40% solution, 3 mL of 20% solution, 3 mL of 10% solution, and 2 mL of 5% solution in a 14 x 89 mm (13.2 mL) thin walled ultra-clear centrifuge tubes (Beckman Coulter, Cat. # 344059). Approximately 250 µl of crude EV sample resuspended in 20 mM Tris-HCl pH 6.0 was slowly dispensed on top of the column and centrifuged at 100,000 x g for 17 h at 4 ⁰C (SW 41 Ti rotor, Beckman Coulter; Optima^TM^ L-90K, Beckman Coulter). After the ultracentrifugation, 4.5 mL solution from the top was discarded and the next 2.1 mL of vesicle containing solution (the layer between 10% and 20% gradient) was collected. Next, the vesicle containing fraction was dispersed in 350 µl fractions in 1.5 mL (9.5 x 38 mm) in 6 polypropylene microfuge tubes (Beckman Coulter, Cat. # 357448) and the volume in each tube was brought up to 1,500 µl using 20 mM Tris-HCl pH 6.0. The tubes were then centrifuged at 100,000 x g for 1 h at 4 ⁰C (TLA 55 rotor, Beckman Coulter; Optima^TM^ Max ultracentrifuge, Beckman Coulter). The pellets from the tubes were pooled together in 500 µl of 20 mM Tris-HCl pH 6.0 buffer and the volume was brought up to 1,500 µl using 20 mM Tris-HCl. It was then centrifuged again at 100,000 x g for 1 h at 4 ⁰C (TLA 55 rotor, Beckman Coulter; Optima^TM^ Max ultracentrifuge, Beckman Coulter). The final pellet containing the purified EV fraction was resuspended in 50 µl of 20 mM Tris-HCl pH 6.0. Samples were kept at 4 ⁰C and used within 48 hours.

### Nano tracking analysis

The analysis of concentration and the size distribution of the sorghum EVs was done using nanoparticle tracking analysis (NTA) as described by Shuler et al. (Shuler et al., 2020). Briefly, a Nanosight NS300 (Malvern Panalytical, Malvern, UK) bearing a 532-nm green laser and NS300 FCTP Gasket (Cat. # NTA4137) was used to determine the size and concentration of the EVs. The parameters used for data acquisition included a camera level of 11, a detection threshold of 6 for the P40, 35 for the density purified, and recording three videos each for 30 seconds. NTA software v3.2 was used for analyzing the data. 30 µl of isolated EVs were added to 970 µl HPLC grade water for P40 and 50 µl of density purified EVs were added to 450 µl of HPLC grade water. The sample was injected into the instrument using a 1 mL sterile BD Plastipak syringe (Becton Dickinson S.A., Madrid, Spain).

### Dynamic light scattering analysis

Dynamic light scattering (DLS) and zeta potential analysis was performed with a Litesizer 500 (Anton Paar, GmbH). For both the P40 and density purified, 30 µl of isolated EVs were resuspended in 970 µl of HPLC water, loaded into a disposable cuvette and the measurements were taken in triplicate.

### Fluorescent labeling and imaging of EVs

EVs isolated in the P40 fraction were stained for 30 minutes at room temperature with 0.5 μM CellTrace ^TM^ Calcein Green (ex488/em521), AM (from ThermoFisher Scientific) and/or 0.5 μM Potomac Gold (ex561/em594) in 20 mM Tris HCL pH 6.0. The stained samples were purified on the iodixanol density gradient as described above. The stained EV samples were mounted on 0.1% poly-L-Lysine coated Cellviz dishes (Cat. # D35-10-1.5N) and the background fluorescence of Potomac Gold was quenched using 50 nM Trypan Blue in HPLC water. Samples were imaged using Borealis Total Internal Reflection Microscopy (B-TIRF) or Spinning disk on an Andor Dragonfly 600 microscope (Oxford Instruments) with a Zyla sCMOS camera and a Leica Plan Apo 63x/1.47 NA oil TIRF objective. A 488 nm excitation laser and a 521/38 nm BP filter was used for Calcein AM. For Potomac Gold, a 561 nm laser and 594/43 nm BP filter was used.

### TEM analysis

400 mesh carbon-coated copper grids were glow discharged on a Pelco easiGlow^TM^ (Ted Pella) prior to incubation with EVs. The samples were negatively stained with 2% aqueous uranyl acetate and imaged using a Zeiss Libra 120 transmission electron microscope operating at 120 kV. Images were acquired using a Gatan Ultrascan 1000 CCD camera.

### Cryo Electron Tomography

Isolation method was stopped at the 40,000 x g spin, and the pellet was resuspended in 50 µl NaHCO3, pH 8.0 and 25 mM trehalose. Samples were then shipped overnight with cold packs to the Brookhaven National Lab. Cryo-EM grids were prepared on a Vitrobot Mark IV and imaged on a Krios G3i Cryo Transmission Electron Microscope operated at 300 kV. Tilt series were acquired using SerialEM or ThermoFisher Tomography with a nominal magnification of 42,000 and a pixel size of 2.2 angstrom on a Gatan K3 camera. Tilt series of +/-50॰ were acquired in 2॰ increments. The energy slit of 20 eV was set on BioQuantum energy filter for tilt series acquisition. Tilt series were aligned and reconstructed using EMAN2. 3D tomograms were loaded into FIJI, average projected and manually analyzed the morphology and size of EVs.

### Correlative imaging by confocal and SEM

Ibidi Grid-50 coverglasses (Cat. # 10817) were prepared using a method adapted from Spakman et al 2018. In short, coverglasses were sonicated for 30 min in 95% isopropanol, rinsed five times with water, sonicated in 5N NaOH for 30 min and rinsed another five times with water. A final 30 min sonication in 100% ethanol and 5 X rinse with water was performed prior to air drying the coverglasses with the grid side faced up. HPLC grade water and ethanol were used for all steps and the glass dish was covered to reduce evaporation during incubations. Cleaned glass was then washed with 100% isopropyl alcohol (IPA) and the excess was baked off at 50 ॰C. The coverglasses were placed grid side faced up on glass stirring rods taped in a large glass dish. The surface of the coverglass was flooded with 0.1 N NaOH and incubated at room temp for 20 min and rinsed 5X with water. Excess water was removed and allowed to air dry. Once completely dry, 5% (3-Aminopropyl)triethoxysilane (APTES) and 95% IPA was applied and incubated for 10 min. The solution was aspirated off and excess was removed with 5X rinses with water. Once completely dry, 1% glutaraldehyde in 0.2 μm filtered PBS pH7.4 was added and incubated for 1 h. Glutaraldehyde solution was removed and 50 μl of the EVs from the P40 fraction pre-labeled with Calcein AM as described above were added to the coverglass and incubated for 30 min. To reduce evaporation, moistened kimwipes were placed in the covered glass dish. 0.2 μm filtered 20 mM Tris-HCl pH 6.0 was added to the coverglasses and then gently removed, care was taken not to let the surface dry. Additional Tris-HCl was added to the coverglass which was then put into an Attofluor Cell Chamber (ThermoFisher, Cat #A7816). A larger area was imaged via spinning disk microscopy by acquiring a 3 x 3 tiled Z stack in 0.01 nm z-interval steps to account for cover glass levelness. Images were stitched in Imaris Stitcher, and Z projections were made using FIJI (Schindelin 2012). After imaging the buffer was carefully removed, and 2% glutaraldehyde in 1 x PBS pH 7.4 was added to and incubated at room temperature for 30 minutes. The coverglasses were then carefully rinsed with HPLC water three times to remove any excess salts for electron microscopy. Additionally, residual immersion oil was removed from the bottom of the coverglass using a kimwipe and sparkle. Samples were then put through an ethanol dehydration series, 25%, 50%, 75%, 95% of HPLC ethanol in HPLC water for 5 minutes each and then finally with 100% anhydrous ethanol for 10 min, then air dried in a covered glass petri dish. The coverglasses were critical point dried using a Tousimis Autosamdri-815B critical point dryer filled with anhydrous ethanol, then mounted on SEM stubs and sputter coated in the ACE600 coater (Leica). SEM image maps were acquired using a ThermoFisher ApreoVS SEM and MAPS version 3.9 software (ThermoFisher Scientific). One or two grid squares were selected based on the confocal image and 7 × 7 image montages were acquired at a setting of 4096X4096 pixel dimension and a 2nm pixel size per image. The image montages were stitched and the confocal images were overlayed and correlated.

### Mass Spectrometry Analysis

EV pellets were denatured in a solution of 8 M urea and 100 mM ammonium bicarbonate, with a final urea concentration of 1 M. To reduce disulfide bonds, samples were mixed with a final concentration of 10 mM Tris (2-carboxyethyl) phosphine hydrochloride (Cat. # C4706, Sigma Aldrich) and incubated at 57°C for 45 min. To alkylate side chains, samples were mixed with a final concentration of 20 mM iodoacetamide (Cat. # I6125, Sigma Aldrich) and incubated at 21°C for 1 hour in the dark. To digest proteins, 500 ng of trypsin (Cat. # V5113, Promega) was added, and the samples were incubated at 37 °C for 14 hours. Following digestion, samples were dried down, resuspended in 0.1% formic acid, desalted using a zip tip (EMD Millipore) and injected into an Orbitrap Fusion Lumos (ThermoFisher) equipped with an Easy NanoLC1200 HPLC (ThermoFisher).

The peptides were separated on a 75 μm x 15 cm Acclaim PepMap100 separating column (Thermo Scientific) downstream of a 2 cm guard column (Thermo Scientific). A 120 min gradient was run from 4% Buffer A (0.1% formic acid in water) to 33% Buffer B (0.1% formic acid in 80% acetonitrile). Peptides were collisionally fragmented using Higher-energy collisional dissociation (HCD) mode, and precursor ions were measured in the Orbitrap with a resolution of 120,000 and a spray voltage set at 1.8 kV. Orbitrap MS1 spectra (AGC 4×10^5^) were acquired from 400-2000 m/z followed by data-dependent HCD MS/MS (collision energy 30%, isolation window of 2 Da) for a three second cycle time. Unassigned and singly charged ions were rejected through the enabling charge state screening. A dynamic exclusion time of one minute was used to discriminate against previously selected ions.

### Database Search and Proteomic Analysis

Proteome Discoverer 2.5 (ThermoFisher Scientific) was used to search the LC-MS/MS data. MS spectra were searched against a *Sorghum bicolor* UniProt database. Parameters for the database search were set as follows: two missed tryptic cleavage sites were allowed for trypsin digested with 10 ppm precursor mass tolerance and 0.05 Da for fragment ion quantification tolerance. Oxidation of methionine was set as a variable modification. Carbamidomethylation (C; +57Da) was set as a fixed modification. Results were filtered using the Percolator node with a FDR of 0.01.

Proteins detected in two out of three samples with a q-value less than 0.01 were selected for further analysis. Signal peptide sequences were predicted using SignalP 6.0 (https://services.healthtech.dtu.dk/services/SignalP-6.0/; Teufel et al., 2022). Transmembrane domains were predicted using TMHMM 2.0 (https://dtu.biolib.com/app/DeepTMHMM/run; Hallgren et al., 2022). Close homologs to Arabidopsis EV proteins (those with an E ≤ 1e-50) were identified using blastp (https://blast.ncbi.nlm.nih.gov/Blast.cgi?PROGRAM<u>=</u> <u>blastp&PAGE_TYPE=BlastSearch&LINK_LOC=blasthome</u>). Gene ontology (GO) enrichment was performed using the Panther Classification System provided by the GO Consortium (http://geneontology.org/). Protein alignments were performed using Clustal Omega (https://www.ebi.ac.uk/Tools/msa/clustalo/; Sievers et al., 2011).

### Protease Protection Assay

The protease protection assay was performed as described in Rutter & Innes (2017). Briefly, EVs were resuspended in 150 mM Tris-HCl pH 7.8 and divided into three samples: A, B and C. Sample A was treated only with Tris-HCl buffer. Sample B was treated with 1μg mL^-1^ trypsin (Promega). Sample C was treated with 1% Triton X-100 (TX100) followed by treatment with 1μg mL^-1^ trypsin (Promega). For the detergent treatment, samples were incubated on ice for 30 min. For the trypsin treatment, samples were incubated in a thermal cycler set at 25 °C for 60 min. All samples were subjected to the same incubation times and temperatures. Following the trypsin treatment step, samples were mixed with 5X SDS buffer and incubated at 95 °C for 5 min.

## Results and discussions

### Isolation of sorghum EVs

Differential centrifugation is the widely used method for isolation of plant EVs from various plant sources. In this study, we used the differential centrifugation method to isolate EVs from sorghum plants by adapting the method described by Rutter and Innes (2017) (Fig. 1). Briefly, ∼14 days old sorghum plants were cut above the soil and vacuum infiltrated in vesicle isolation buffer (VIB). The apoplastic wash (AW) was collected from the infiltrated plants at a low speed of 700 x g. Initially, we continued with only the AW for the downstream differential centrifugation steps. However, the low yield of EVs from our multiple trials prompted us to check if EVs were getting released into the VIB by the repetitive changes in vacuum pressure. To look for EV-like particles in the residual VIB from vacuum treatments (vAW), we stained the vAW with the membrane dye Potomac Gold. Spinning disk confocal microscopy detected EV-like objects in vAW (Fig. S1). Therefore, we used the combined AW and vAW for the subsequent downstream centrifugation steps. The P40 sorghum EVs were obtained by centrifuging the combined AW and vAW initially at 10,000 x g to remove the large cellular debris followed by two rounds of subsequent centrifugation at 39,800 x g (Fig. 1). Further purification of EVs was done using discontinuous iodixanol (OptiPrep) gradient (Rutter and Innes, 2017). Briefly, the P40 pellet was added on the gradient column of iodixanol and centrifuged for 17 hours at 100,000 x g. The interlayer fractions between 10% and 20% gradient were collected and centrifuged again at 100,000 x g for 1 h to obtain the purified EV fractions.

**Figure 1-.**
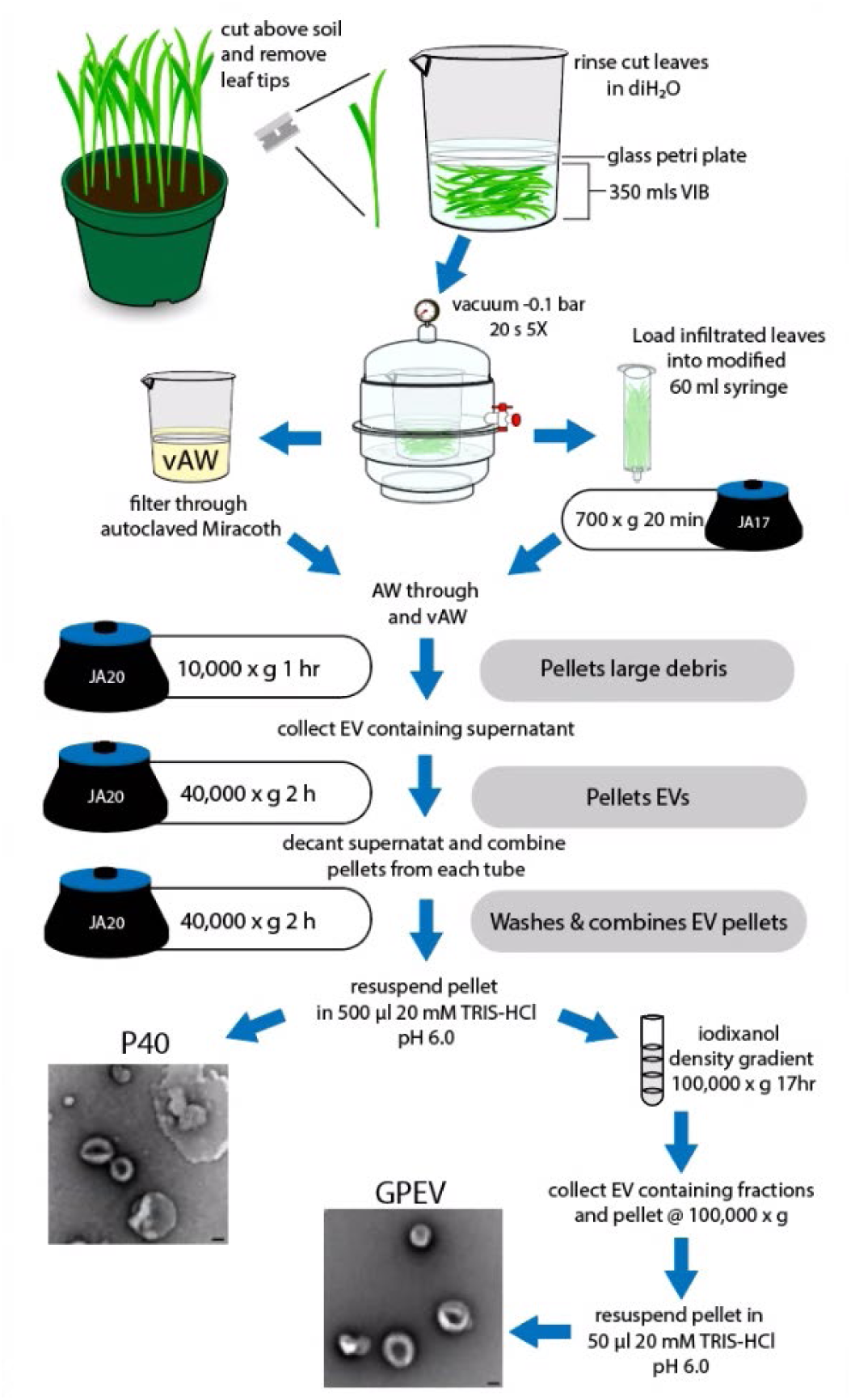
Sorghum EV isolation protocol.

### Characterization of sorghum EVs by particle analysis

We used nanoparticle tracking analysis (NTA) to determine the size and yield of the particles in the P40 pellet. The EV isolation was divided into two portions, the P40 containing pellet and density purified EVs. The P40 EVs had a mean size of 170.5 nm and density purified vesicles had a mean size of 193.3 nm. The observation of multiple peaks and large standard deviation in the NTA analysis indicates a heterogeneous population of EV particles of varying sizes. Similar size distribution was reported for EV-like particles from Arabidopsis leaves that were obtained from the P40 (Rutter and Innes, 2017). They demonstrated that the particle size of the P40, isolated from the Arabidopsis leaves using similar techniques as described in our study, ranged approximately from 50 to 300 nm in diameter; particles around 150 nm in diameter were the predominant species (Rutter and Innes, 2017). NTA has a detection size limit of 1000 nm, and therefore, dynamic light scattering (DLS) was used to detect larger vesicles. DLS of P40 EVs has a hydrodynamic radius mean of 1,517.6 nm with a standard deviation of 1619 and two peaks at 160.88 and 1521.07 nm. A large amount of variation was observed in the peaks and percent distribution of the technical replicates. In contrast, the density purified EVs had a single peak with a mean value of 182.23 nm with a relatively low standard deviation of 9.85 nm, suggesting the EV size is more homogeneous.

### Characterization of morphology

We first analyzed the morphology of the EV-like particles obtained from sorghum leaves by TEM, which showed numerous cup-shaped objects of various sizes (Fig. 2a,b). The appearance of our EVs as cup-shaped is similar to TEM images of EVs obtained from Arabidopsis leaves through differential centrifugation and PDV isolated from strawberry, lemon, orange, and cabbage juices (Rutter and Innes, 2017; Liu et al., 2020b; Baldini et al., 2018; Berger et al., 2020; Yang et al., 2020; Perut et al., 2021; You et al., 2021). TEM images of the P40 fraction revealed a heterogeneous size distribution of EV-like particles. In contrast, the density purified EVs were more homogeneously-sized. These differences in size distribution seen by TEM are in agreement with the NTA and DLS measurements.

**Figure 2-.**
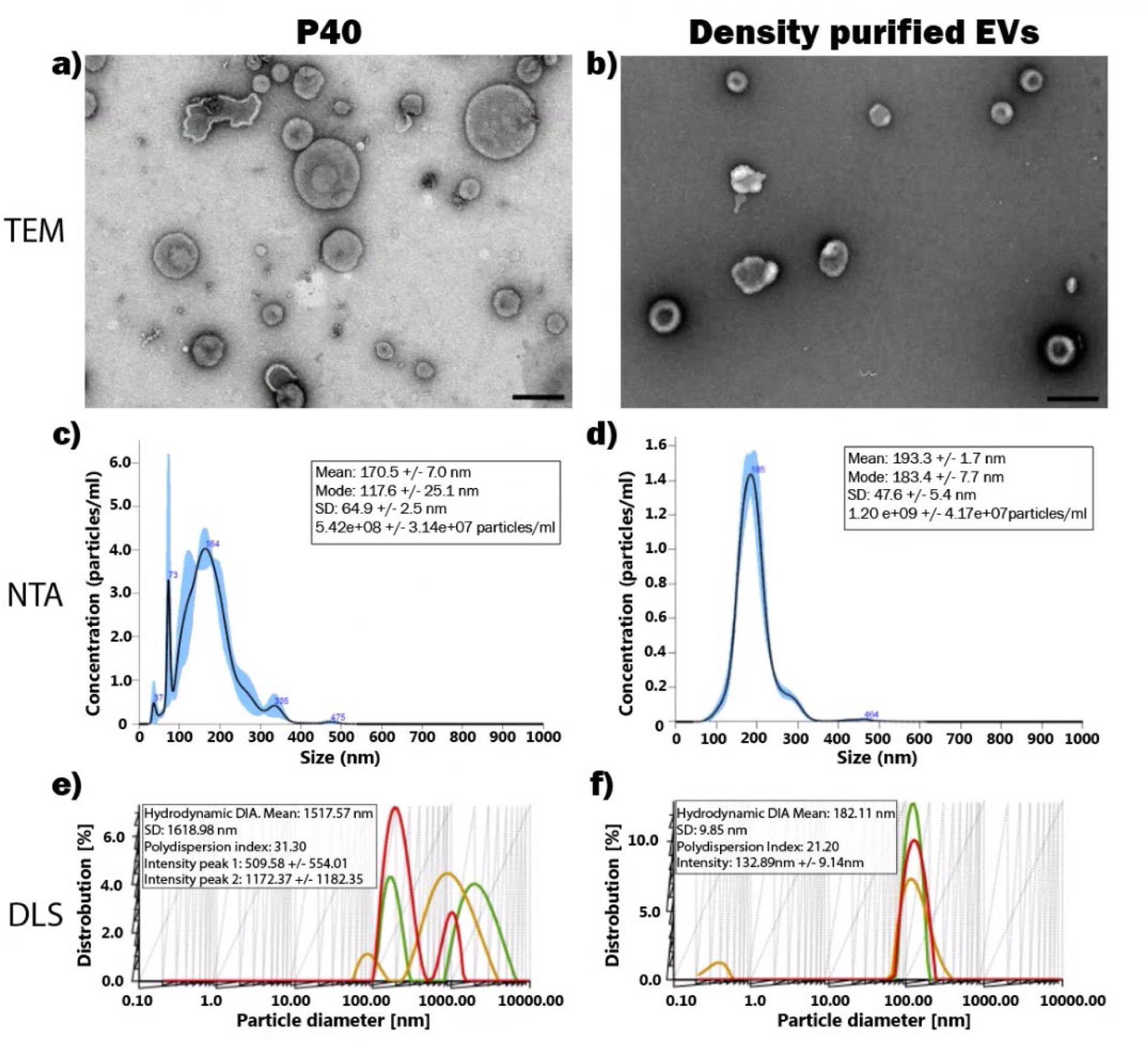
Characterization of P40 and density purified EVs. (a,b) P40 (a) and iodixanol density purified EVs (b) were negatively stained and imaged by TEM. Scale bar 200 nm. (e,f) Nanotracking analysis (NTA) of EVs. Light blue represents the standard error of the mean of 3 technical replicates. (c,d) Dynamic light scattering (DLS) of EVs. Colors represent technical replicates.

To further characterize the EVs, we performed Cryo-Electron Tomography (cryo-ET). The P40 fraction with greater variation was chosen for cryo-ET to obtain a more complete analysis of types of EV-like particles that can be purified from Sorghum. The cryo-ET images analysis of 483 vesicles showed a variety of vesicles that were split into 2 distinct categories: single layer (444) and multi-layer (39) (Figure 3). EVs in the single layer category were vesicles with one lipid bilayer (Fig 3a) and had a mean diameter of 135.58 nm and a median diameter of 104.03 with a standard error of 5.78nm. The multi-layer EVs (Fig 3b) had more than one lipid bilayer with the most common being two bilayers as seen in Fig. 3c. These EVs were larger with a mean diameter of 322.31nm and a median diameter of 274.67 nm with a standard error of 31.24nm. We also quantitated the number of EVs that appeared electron dense in the cryo-ET micrographs (Fig 3c).

**Figure 3-.**
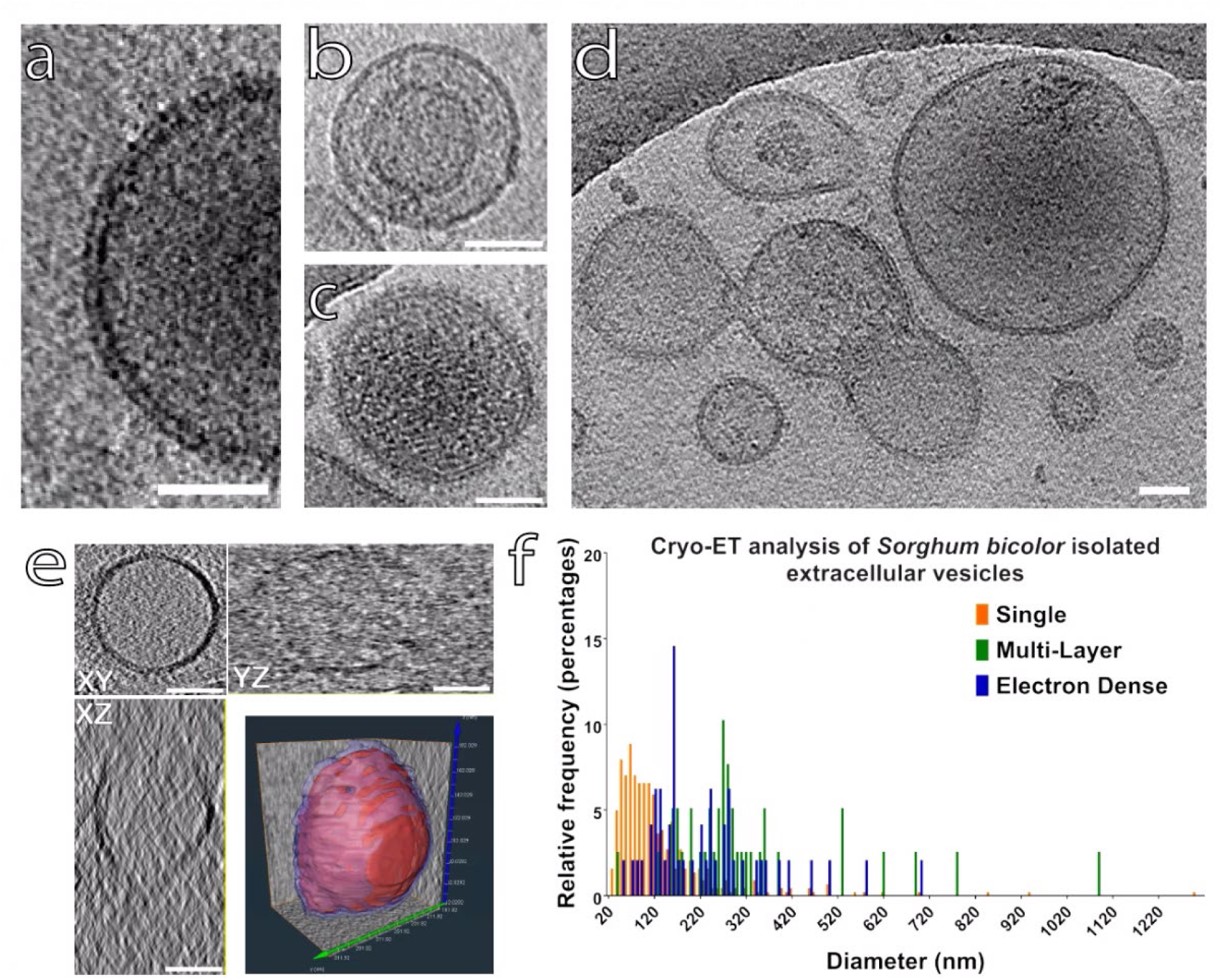
Analysis of cryo-ET sorghum P40 EVs. a) Clear visualization of the lipid bilayer b) Multi-layer c) Electron dense vesicle d) field showing the mixed varieties seen in the P40 pellet e) Single layer vesicle that was rendered in Amira. The red core shows the inner boundary of the vesicle and the light blue are the fringes on the vesicle. f) A histogram of the measured vesicles split into 3 categories. The mean size is 150.95 nm and a median of 110.00 nm. Scale bar is 50 nm.

Forty-three EVs (8%) were electron dense, which suggests these have a higher concentration of cargo. Both single and multi-layered EVs appeared electron dense and a difference in density between the different layers of multi-layered EVs was observed (Supplementary Fig. 2). Another prominent characteristic of these EVs were structures coating or protruding from the surface (Fig. 3c). Additionally some images contained vesicles that appear to be connected to each other (Fig 3d). 3D rendering of a single vesicle showed that they were relatively spherical in structure, and it was possible to separate the fringed edge from the more defined lipid bilayer (Fig 3e). Figure 3f shows the distribution of the three categories of manually quantified EVs.

### Confocal microscopy of sorghum EVs

To characterize sorghum EVs using light microscopy approaches, we developed a method that uses both a soluble and lipophilic membrane fluorescent dye. First, we explored the soluble, membrane permeable fluorescent dye, Calcein AM, for labeling isolated sorghum EVs. Calcein AM, the acetomethoxy derivative of calcein, is commonly used for cell viability assay, but has also been used to label EVs (Hiraoka and Kimbara, 2002; Bauer et al., 2003; Bratosin et al., 2005; Miles et al., 2015 Gray et al. 2015). Calcein AM green is a non-fluorescent esterase-dependent substrate and lipophilic in nature, which allows it to passively cross membranes (Bratosin et al., 2005). Once inside EVs, calcein AM is hydrolyzed to become fluorescent and EV membrane impermeable. After incubating the P40 sorghum EVs with calcein green, we observed small fluorescent foci using fluorescence light microscopy (Fig. 4). To determine if these spots corresponded to EVs, we conducted correlative light and electron microscopy (CLEM) (Fig. 5). EVs were deposited on gridded coverglasses and imaged using the Dragonfly spinning disk and subsequently by scanning electron microscopy. 122 stained vesicles corresponded to round vesicles of the expected size range between the confocal and SEM images.

**Figure 4-.**
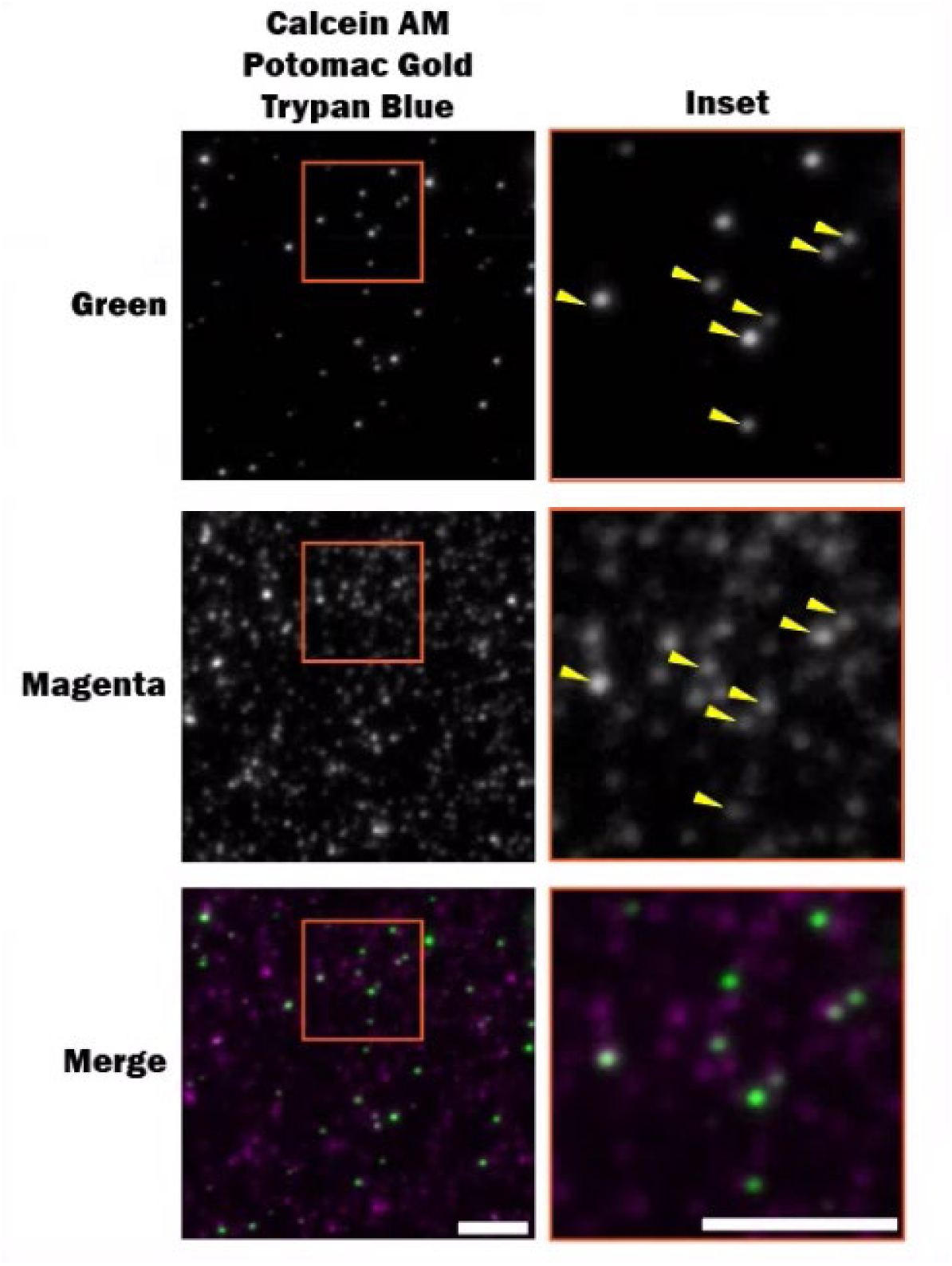
bTIRF imaging of sorghum density purified EVs. Images displayed with a median filter of 1. Scale bar 5 μm.

**Figure 5-.**
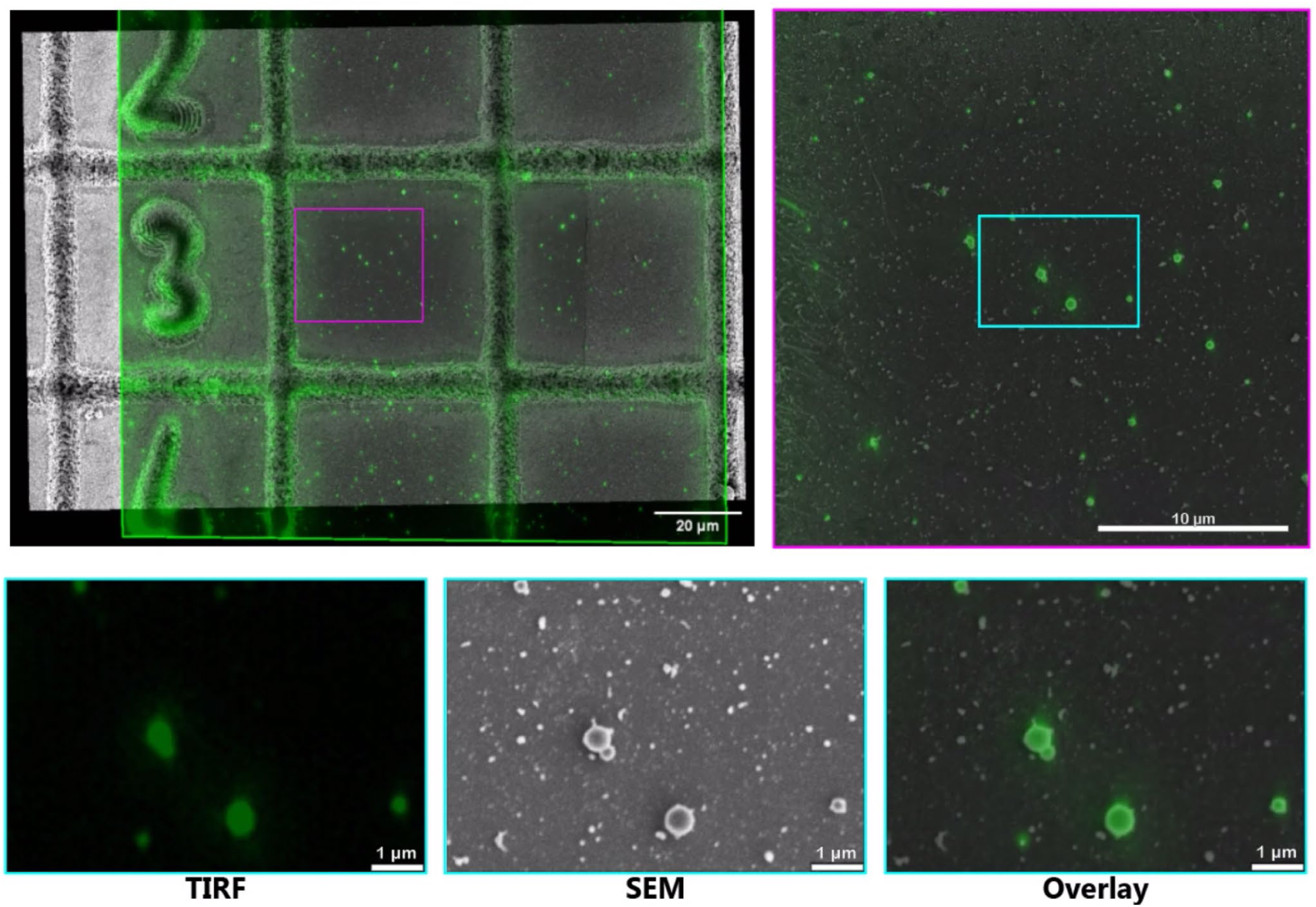
Correlative image analysis. Spinning disk image of the calcein AM stained sorghum vesicles overlaid on the SEM data.

To determine if these fluorescent foci were membrane bound EVs, we also labeled the EVs with a lipophilic membrane fluorescent dye and looked for co-localization between the two fluorescent signals. Our initial experiments of EV staining with PKH26 were not successful (data not shown), and therefore, we evaluated Potomac Gold, a non-specific membrane dye known to label membrane bound organelles in bacteria and mammalian cells (Spahn et al., 2018). Fig. 4 shows colocalization of both the calcein AM green and Potomac Gold-stained objects indicating that the calcein AM green labeled vesicles are membrane bound EV-like particles.

### Proteomic Analysis

To better understand the protein content of sorghum EVs, as well as identify potential EV markers, we analyzed sorghum vesicles using liquid chromatography with tandem mass spectrometry (LC-MS/MS). To remove any contaminating, non-vesicular particles, the crude EV pellets were first purified on a discontinuous iodixanol gradient, as described in Rutter and Innes (2017). Analysis of three biological replicates of purified EVs yielded 454 proteins. From this list, we selected proteins for further analysis that were detected in two out of three replicates with a q-value ≤ 0.01. The final sorghum EV proteome consisted of 437 proteins.

As membranous compartments involved in the unconventional secretion of proteins, EV proteomes contain a higher proportion of membrane proteins and proteins lacking a N-terminal signal peptide (SP) for classical secretion (Rosa-Fernandes et al., 2017). Online software predicted that 14.87% (65/437) of the sorghum EV proteins possessed a SP (Fig. 6A) (Teufel et al., 2022), while 37.30% (163/437) of EV proteins were predicted to have at least one transmembrane region (TMR) (Fig. 6B) (Hallgren et al., 2022). This agrees well with data on animal EVs proteomes. Around 80% of the top 100 most identified animal EV proteins lack a SP, and 40% possess at least one TMR (Rosa-Fernandes et al., 2017). Similarly, 84% of proteins associated with Arabidopsis EVs lacked a SP and 34% possessed at least one TMR (Rutter and Innes, 2017).

**Figure 6-.**
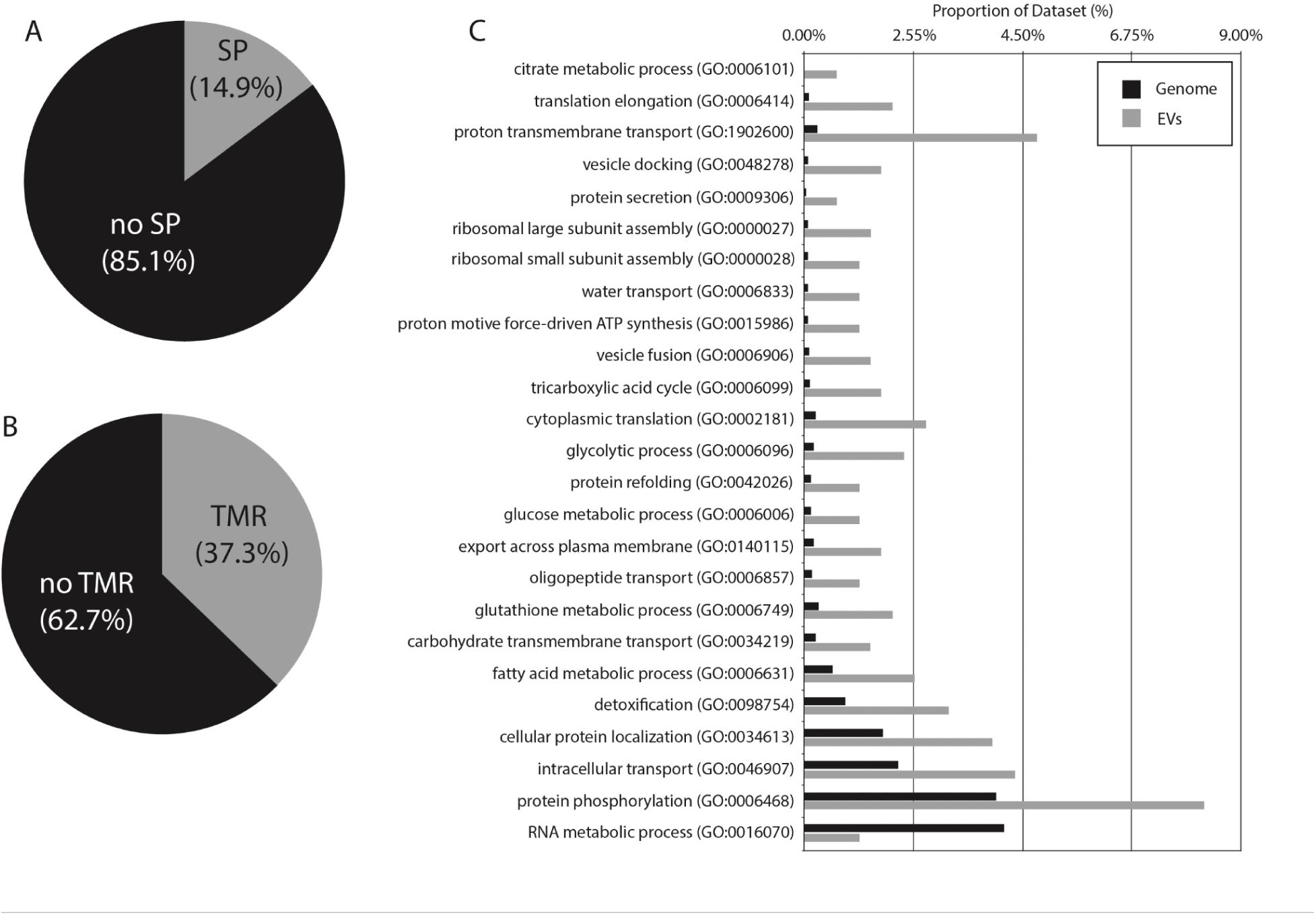
Sorghum EV Proteome. Online software predicts 14.9% of Sorghum EV proteins possess a signal peptide (SP) (A). Online software predicts 37.3% of Sorghum EV proteins possess at least one transmembrane region (TMR) (B). Gene ontology (GO) term enrichment analysis revealed both enriched and depleted terms in the Sorghum EV proteome (C).

Gene ontology (GO) term enrichment analysis suggests that the sorghum EV proteome is enriched for proteins involved in vesicle docking/fusion, translation/ribosome, transmembrane transport, carbohydrate/amino acid/lipid metabolism and protein folding (Fig. 6C). Similar GO terms were enriched in EVs from Arabidopsis (Rutter and Innes, 2017; He et al., 2021). However, approximately 32% of the sorghum EV proteome was annotated as “uncharacterized,” making it difficult to gain a thorough understanding of the proteome and predict which functions the EVs may possess.

To gain deeper insight into the sorghum EV proteome and search for similarities between monocot and dicot EVs, we took the list of Arabidopsis EV proteome published in Rutter and Innes (2017) and searched for the closest homologs (E value ≤ 1e-50) present in the sorghum EV proteome. We found that 60% of Arabidopsis EV proteins had a homolog in the sorghum EV proteome (Fig. 7). This included homologs to the marker proteins PEN1 (AT3G11820.1), PATELLIN 1 (PATL1, AT1G72150.1), the ABC transporter PENETRATION3 (PEN3,AT1G59870.1) and TET8 (AT2G23810.1), as well as the proposed RNA-binding annexin proteins ANN1 (AT1G35720.1) and ANN2 (AT5G65020.1) (Fig. 7). Biological function GO term enrichment of sorghum homologs suggests a core of proteins enriched for vesicle docking/fusion, transport across membranes, exocytosis, carbohydrate metabolism and translation. We also identified several homologs to Arabidopsis stress-and defense-related proteins, suggesting that sorghum EVs may also have a role in immunity (Table. S1).

**Figure 7-.**
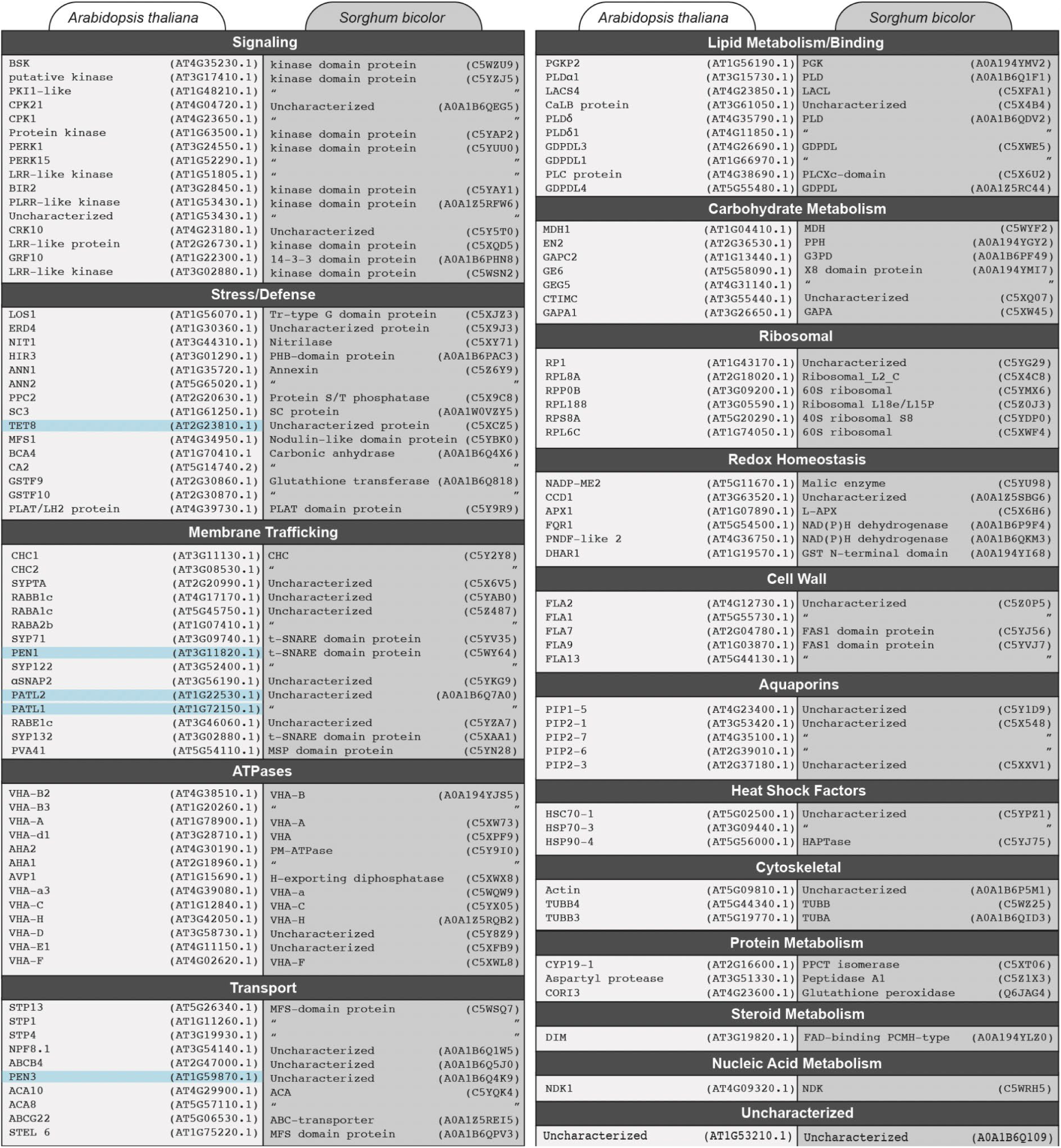
Sorghum homologs to *Arabidopsis* EV proteins. *Arabidopsis* EV proteins were analyzed using NCBI’s blastp to identify the closest homolog (E ≤ 1e-50) present in the Sorghum EV proteome. Proteins were categorized roughly according to biological function. Proteins commonly identified as Arabidopsis EV markers are highlighted in blue. The presence of a quotation mark indicates that the closest Sorghum homolog is the same as the one listed immediately above.

To verify our proteomic data and establish tentative EV markers, we conducted a protease protection assay and probed the treated EVs using antibodies raised against common Arabidopsis EV markers PATL1, PEN1 and TET8. In this assay, an EV pellet is divided into three samples and treated with either buffer, trypsin protease or Triton X-100 (TX100) detergent followed by trypsin. Proteins protected within the lumen of EVs will remain intact in the presence of trypsin, unless the EV membranes are first solubilized using TX100. When using antibodies specific for Arabidopsis PATL1 or PEN1, we detected bands in buffer and trypsin-treated EV samples (Fig. 8). These bands disappeared when the EVs were pre-treated with detergent. The Arabidopsis TET8 antibody was able to detect a strong band of the expected size in lysate samples, but was unable to detect any bands in EV samples. The results suggest that sorghum homologs of Arabidopsis PEN1 and PATL1 are similar enough to be bound by Arabidopsis-specific antibodies and that these homologs are protected within the lumen of membranous compartments. The signal for the band cross-reacting with the PEN1 antibody was particularly strong in both lysate and EV samples. Because sorghum possesses only a single close homolog to PEN1, the t-SNARE C5WY64, the band we detected likely represents the true sorghum PEN1 homolog (Fig. S4). Combined, our data suggests that PEN1 may serve as a useful marker for the identification of both Arabidopsis and sorghum EVs.

**Figure 8-.**
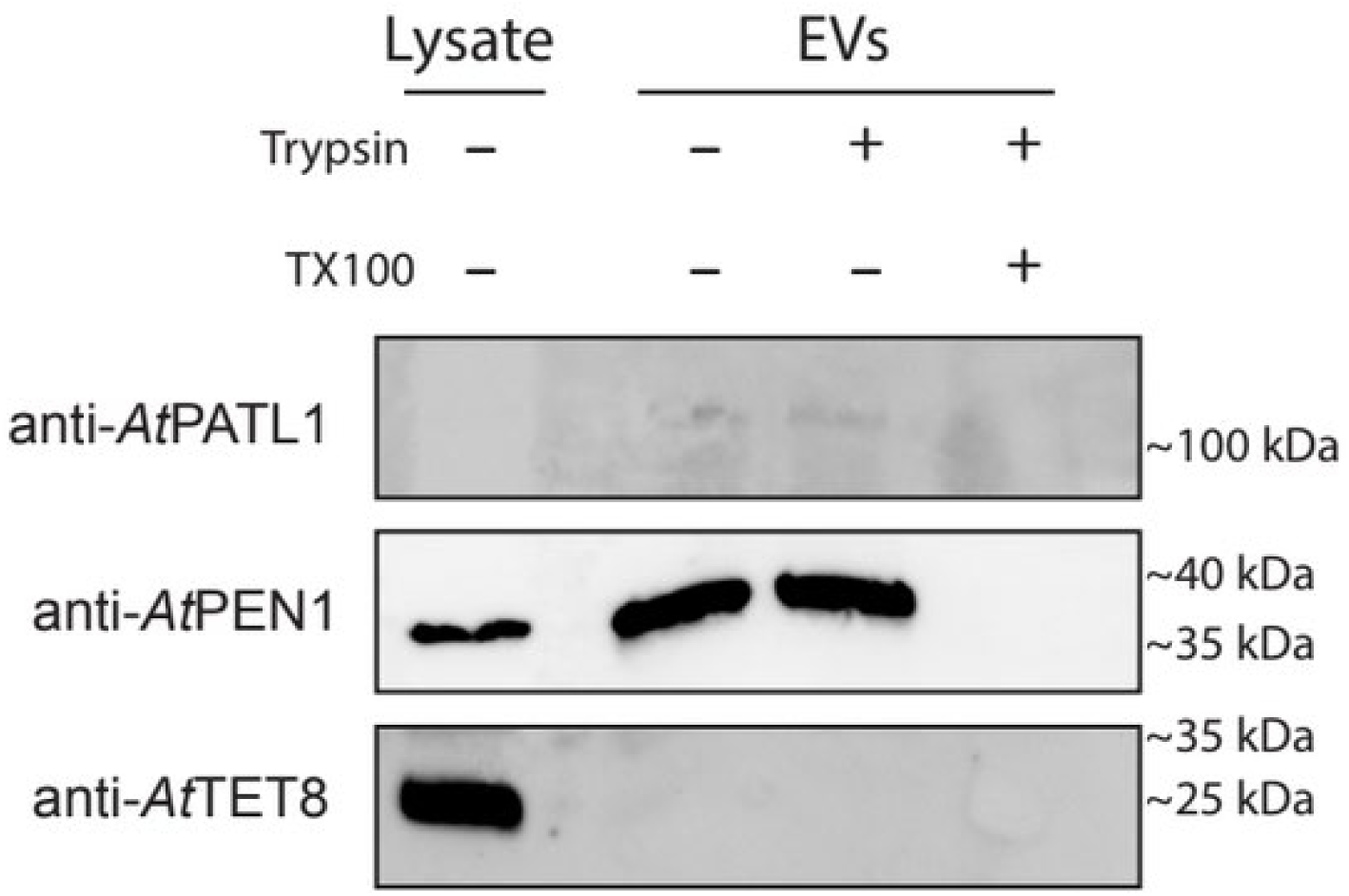
Protease protection assay. Sorghum EVs were treated with buffer, trypsin and trypsin + Triton X-100followed by trypsin. Samples were then probed using antibodies raised against Arabidopsis EV marker proteins.

## Conclusion

Considering research into plant EVs is still in its infancy, not much is known about the proteins associated with EVs from various plant species. Evidence suggests that EVs from Arabidopsis and sunflower are enriched for proteins involved in stress responses and defense, but relatively nothing is known about the protein content of EVs produced by monocots (Rutter & Innes, 2017; Regente et al., 2017). The syntaxin protein PENETRATION 1 (PEN1) and TETRASPANIN 8 (TET8) have been identified as useful marker proteins for the identification and tracking of EVs in Arabidopsis (Rutter & Innes, 2017; Cai et al., 2018), but it is uncertain if homologous proteins can be used to mark monocot EVs. The lack of an appropriate EV marker was a limitation for further studies. In this study, we described an improved and optimized protocol involving differential centrifugation for the isolation of EVs from sorghum leaves and have identified EV markers by mass spectrometry. This proteomic analysis shows that a majority of proteins in the Arabidopsis EV proteome have a close homolog in the sorghum EV proteome, including the commonly used EV markers, PEN1, PATL1 and TET8. Our study further establishes PEN1 and PATL1 as universal plant EV markers, and when combined with our isolation protocol, it lays the foundation for future studies in sorghum and other monocots.

A major aim of this study was to adapt the Rutter et. al (Rutter, 2017) Arabidopsis EV isolation protocol for the monocot, sorghum. In general monocot leaves are more hydrophobic than Arabidopsis leaves, and a critical addition to the sorghum protocol was the snipping of the leaf tips to assist with the vacuum infiltration of the vesicle isolation buffer. The second major change was the retention of the buffer used for vacuum infiltration apoplastic wash (vAW) for subsequent EV isolation. We demonstrated that EVs get released into the buffers during vacuum infiltration, and the yield of EVs can be improved by using both cAW and vAW for downstream processes after infiltration. The addition of the vAW to the EV isolation protocol allowed us to capture vesicles released from the leaves caused by the pressure changes during the vacuum infiltration. The pressure changes cause VIB to enter and exit the apoplastic space via stomates and the cut ends of the vasculature at the ends of the leaves. This process mimics a “washing” of VIB in and out of the leaf with the eventual replacement of all the apoplastic air with VIB, generating a vacuum-based apoplastic wash that we call vAW. EV isolations from the eudicot, spinach, had a significant amount of EVs in the vAW (data not shown), and this suggests EVs in the vAW is not specific to sorghum or monocots. Future studies in other systems, such as Arabidopsis, would help determine if EV release into vAW is a common outcome.

Cryo-ET analysis of grapefruit-derived nano-particles extracellular vesicles showed similar single and double-layered vesicles (Garavea et al., 2021) Many of the vesicles also showed increased electron density, indicating the presence of cargo. Structural analysis of the sorghum EVs revealed a diverse population of vesicles that were electron dense, multi-layered and a median size of 110 nm. Similar morphological diversity is seen in mammalian derived vesicles. (Emelyanov 2020, Matthies 2020)

We also explored appropriate dye for labeling EVs and described that calcein AM green and Potomac Gold are suitable fluorescent dyes for labeling EVs for confocal microscopy. From these experiments we showed that calcein AM robustly labeled these vesicles.

In the absence of a suitable EV marker, we attempted to stain Sorghum EVs using fluorescent dyes. The dyes that typically have been reported to label vesicles from various plant sources are mainly the lipophilic membrane staining dyes. It was reported that PKH26 and PKH67, the commonly used lipophilic membrane staining dye, were used to label PDVs isolated from various plant sources. For example, PKH26 were reported to be used for labeling PDVs from strawberry, grape, grapefruit, ginger, carrot, and lemon juices whereas PKH67 was used to label PDVs from cabbage (Mu et al., 2014; Raimondo et al., 2015; Perut et al., 2021; You et al., 2021). Another study reported using Dil (Dioctadecyl-3,3,3,3-tetramethylindodicarbocyanine), another lipophilic membrane dye, to label PDVs isolated from lemon juice (Yang et al., 2020). Cell mask orange (CMO) is also a commonly used lipophilic membrane binding dye that was reported to label EVs isolated from *Arabidopsis thaliana* (Liu et al., 2020b). These dyes have long aliphatic tails that intercalate into membranes, which could potentially change the integrity of the membrane, causing leakiness (Fitzner 2011). Furthermore, a detailed study on the effect of PKH family of dyes on EVs showed they increase their size. In contrast, luminal dyes, such as CFSE do not increase the size of EVs, although they will only fluoresce in EVs that can hydrolyze esters and label only a subset of EVs (Dehghani, M 2020). Here, we used both a luminal, ester-based dye, calcein AM, and a new lipophilic dye Potomac Gold (Spahn, C. K. 2018). Unlike PKH dyes, Potomac Gold does not have a long aliphatic chain and its mechanism of labeling is unknown. It is unlikely to just bind to the surface of EVs like FM4-64, which has been used previously to stain EVs from sunflowers (Regente et al., 2017), because the addition of trypan blue quenches the fluorescence of Potomac Gold bound to the surface of the coverglass, but not from the EVs. This suggests that the lipid bilayer is protecting Potomac Gold from the trypan blue. Lipophilic dyes can also dissociate from membranes, making them unreliable for tracking vesicles (Munter et al., 2018).

Proteins associated with Sorghum EVs were similar to those found in other plant EV proteomes. A small percentage of proteins possessed a SPs, and a larger percentage possessed at least one TMR. Sorghum EVs were enriched for proteins involved in vesicle trafficking, the ribosome, transmembrane transport, general metabolism and protein folding. Similar biological functions were identified for EV proteins from Arabidopsis (Rutter and Innes, 2017; He et al., 2021). Moreover, blastp analysis identified close homologs for 60% of Arabidopsis EV proteins in Sorghum EVs. Combined, the data suggests that the protein content of plant EVs is largely conserved between monocots and dicots. The pathways of EV biogenesis and ultimate function of these vesicles may be similarly conserved. The similarity of monocot and dicot EV proteins suggests that established EV markers for Arabidopsis could be used to identify and validate EVs from Sorghum. To this end, we were able to show that antibodies raised against native Arabidopsis EV marker proteins, especially PEN1, were able to detect homologous proteins in Sorghum EVs. Although we were able to detect bands of an expected size using immunoblots, more work is required to confirm detection of a legitimate homolog. PEN1 belongs to the SYP12 clade of syntaxins that has a conserved function in fungal defense responses. PEN1 does not play a direct fungal pathogenesis, but rather, it is required for the delivery of defense related compounds (Liu M 2022). Similarly, PEN1 is more likely involved in EV biogenesis and not a functional cargo. In general, what is considered the cargo of an EV proteome can vary. In this study, we are identifying all proteins that are enriched during a sorghum EV isolation. This includes proteins inside the EVs, its lipid bilayer, the outside surface of the EV, and even proteins that co-enrich during the isolation.

## Acknowledgements

We would like to thank Dr. Jonathan Trinidad and the Laboratory for Biological Mass Spectrometry at Indiana University for their help generating the proteomic data and Shannon Moda at the University of Delaware Bio-Imaging Center for assistance with transmission electron microscopy, We would also like to thank Luke Lavis and Janelia for the Potomac Gold dye. Funding for staff and reagents was provided by the DOE (DE-SC0020348). Microscopy equipment was acquired by shared instrumentation grants (S10 OD030321 & S10 OD025165) and microscopy access was supported by grants from the NIH-NIGMS (P20 GM103446 & P20 GM139760) and the State of Delaware. The Laboratory for BioMolecular Structure (LBMS) is supported by the DOE Office of Biological and Environmental Research (KP1607011).

