## Supplemental Figures for "Characterization of extracellular vesicles isolated from *Sorghum bicolor* reveals a conservation between monocot and eudicot extracellular vesicle proteomes"

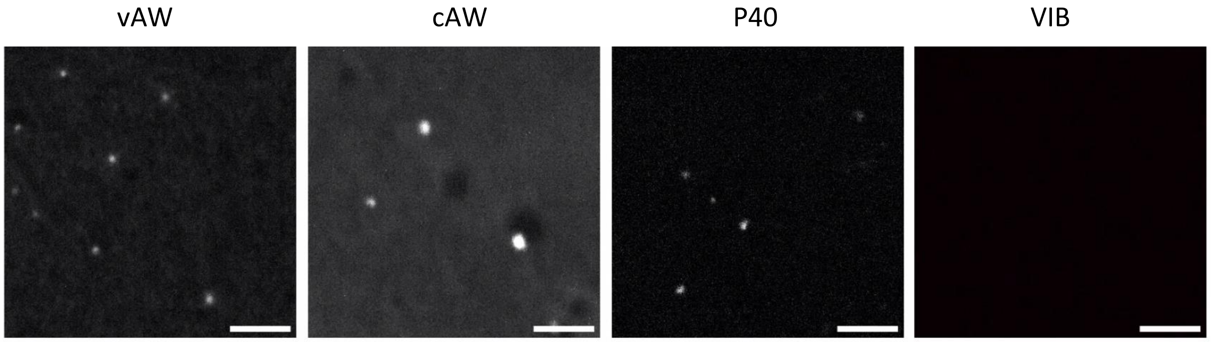


**S.1** Vacuum-based apoplastic wash (vAW), centrifugation-based apoplastic wash (cAW), P40 fraction and vesicle isolation buffer (VIB) negative control all stained with Potomac Gold and imaged via spinning disk confocal microscopy. Scale bar 10 μm.

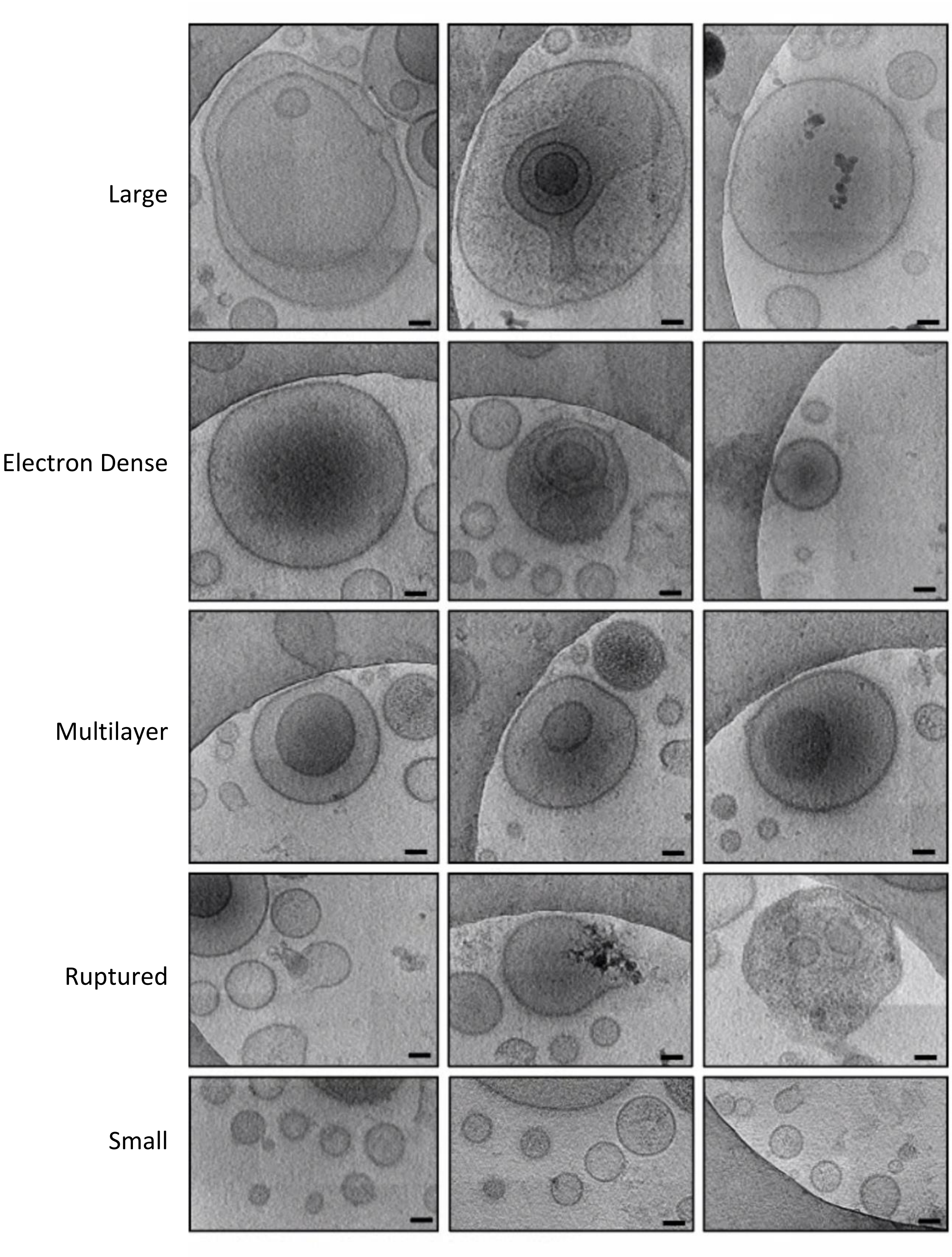


**S2:** variation of vesicles seen in cryo-ET images. Many of the electron dense vesicles also show a hazy fringe around the outside of the lipid membrane. Scale bar = 50 nm

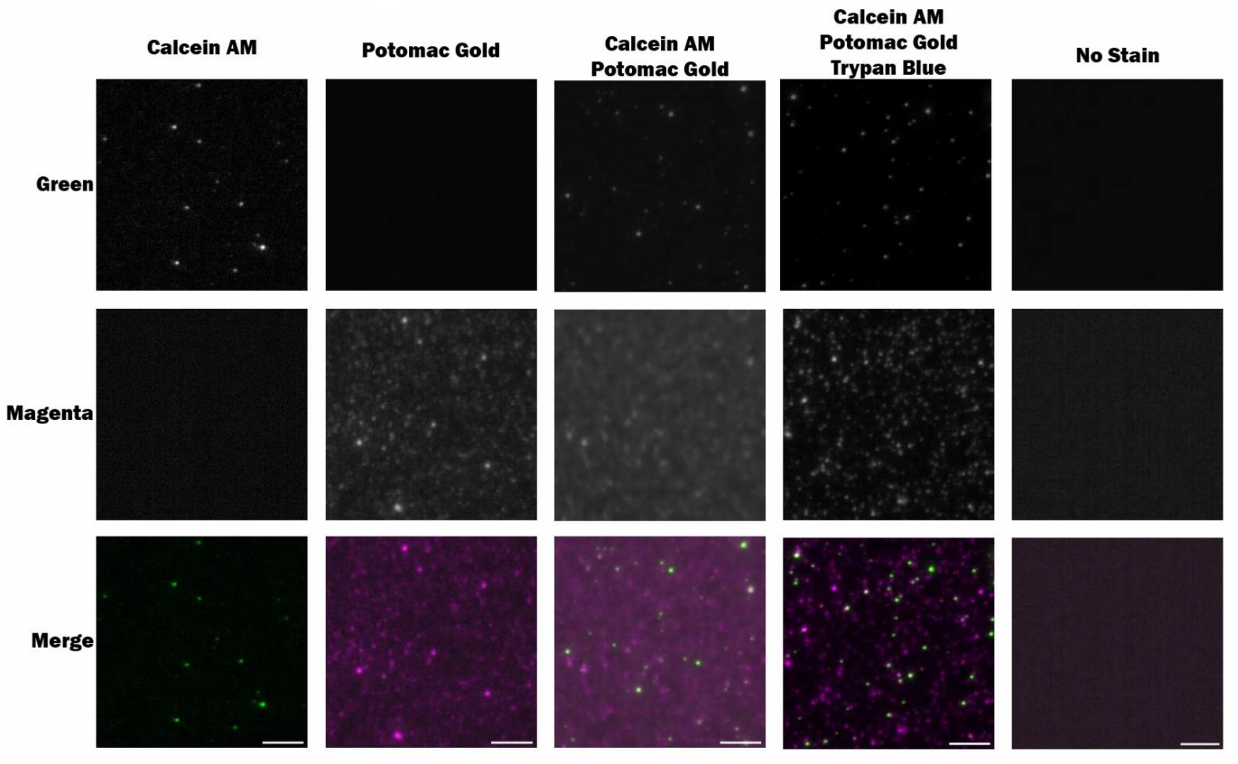


**S3:** Density purified vesicles stained with Calcein AM, Potomac Gold, both stains combined, both stains with the addition of 50 nM trypan blue and a buffer stain control. Scale bar equals 5 μm.

```
AtPEN1 (AT3G11820)   MNDLFSSSFSRFRSGEPSPRRDVAGGGDGVQMANPAGSTGGVNLDKFFEDVESVKEELKE   60
SbPEN1 (C5WY64)      MNSLFSSSWKRGGGDDG------DIESGSVEMSAPPGAAAGASLDRFFEDVESIKDELRD   54
                     **.*****:.*  ..:          ...*:*: * *::.*..**:*******:*:**::

AtPEN1 (AT3G11820)   LDRLNETLSSCHEQSKTLHNAKAVKDLRSKMDGDVGVALKKAKMIKVKLEALDRANAANR   120
SbPEN1 (C5WY64)      LERIQRSLHDGNEAGKSLHDASAVRDLRARMDADVSAAIKKAKVVKLRLESLDRANAANR   114
                     *:*::.:* . :* .*:**:*.**:***::**.**..*:****::*::**:*********

AtPEN1 (AT3G11820)   SLPGCGPGSSSDRTRTSVLNGLRKKLMDSMDSFNRLRELISSEYRETVQRRYFTVTGENP   180
SbPEN1 (C5WY64)      SVPGCGPGSSTDRTRTSVVAGLRKKLRDSMESFSSLRSRVASEYRDTVARRYFTVTGTQP   174
                     *:********:*******: ****** ***:**. **. ::****:** ******** :*

AtPEN1 (AT3G11820)   DERTLDRLISTGESERFLQKAIQEQGRGRVLDTINEIQERHDAVKDIEKNLRELHQVFLD   240
SbPEN1 (C5WY64)      DEATLDALAESGEGERFLQRAIAEQGRGQVLGVVAEIQERHGAVAELERSLLELQQVFND   234
                     ** *** * .:**.*****:** *****:**..: ******.** ::*:.* **:*** *

AtPEN1 (AT3G11820)   MAVLVEHQGAQLDDIESHVGRASSFIRGGTDQLQTARVYQKNTRKWTCIAIIILIIIITV   300
SbPEN1 (C5WY64)      MAVLVAAQGEQLDDIEGNVGRARSFVDRGREQLQVARKHQKSTRKWTCIAILILLVIILV   294
                     *****  ** ******.:**** **:  * :***.** :**.*********:**::** *

AtPEN1 (AT3G11820)   VVLAVLKPWNNSSGGGGGGGGGGTTGGSQPNSGTPPNPPQARRLLR   346
SbPEN1 (C5WY64)      IVLPIVLQNTKKN---------------------------------   307
                     :** ::   .:..
```

**S4:** Clustal alignment of Arabidopsis PEN1 (AT3G11820) and the sorghum homolog C5WY64.
